# Cigarette smoking increases coffee consumption: findings from a Mendelian randomisation analysis

**DOI:** 10.1101/107037

**Authors:** Johan H Bjørngaard, Ask Tybjærg Nordestgaard, Amy E Taylor, Jorien L Treur, Maiken E. Gabrielsen, Marcus R Munafò, Børge Grønne Nordestgaard, Bjørn Olav Åsvold, Pål Romundstad, George Davey Smith

## Abstract

**Background:** Smokers tend to consume more coffee than non-smokers and there is evidence for a positive relationship between cigarette and coffee consumption in smokers. Cigarette smoke increases the metabolism of caffeine, so this association may represent a causal effect of smoking on caffeine intake.

**Methods:** We performed a Mendelian randomisation analysis in 114,029 individuals from the UK Biobank, 56,664 from the Norwegian HUNT study and 78,650 from the Copenhagen General Population Study. We used a genetic variant in the *CHRNA5* nicotinic receptor (rs16969968) as a proxy for smoking heaviness. Coffee and tea consumption were self-reported. Analyses were conducted using linear regression and meta-analysed across studies.

**Results:** Each additional cigarette per day consumed by current smokers was associated with higher coffee consumption (0.10 cups per day, 95% CI:0.03,0.17). There was weak evidence for an increase in tea consumption per additional cigarette smoked per day (0.04 cups per day, 95% CI:-0.002,0.07). There was strong evidence that each additional copy of the minor allele of rs16969968 (which increases daily cigarette consumption) in current smokers was associated with higher coffee consumption (0.15 cups per day, 95% CI:0.11,0.20), but only weak evidence for an association with tea consumption (0.04 cups per day, 95% CI:- 0.01,0.09). There was no clear evidence that rs16969968 was associated with coffee or tea consumption in never or former smokers.

**Conclusion:** These findings suggest that higher cigarette consumption causally increases coffee intake. This is consistent with faster metabolism of caffeine by smokers, but may also reflect behavioural links between smoking and coffee.

## Introduction

Smoking and coffee consumption have been shown to be strongly positively associated; smokers consume more coffee than non-smokers and, within smokers, smoking heaviness is associated with higher coffee consumption (1). Whether this relationship is specific to coffee, or extends to other caffeinated beverages, is less clear; however, a recent analysis in a UK cohort found a similar positive relationship with tea, suggesting it could apply more generally to caffeine (2).

Given the known harmful effects of smoking and the widespread interest in the health effects of coffee or caffeine, which have been implicated (often as a protective factor) in a number of diseases including cancer, depression and diabetes (3–7), it is important to understand what drives this association. Cigarette smoking induces the cytochrome P450 1A2 (*CYP1A2*) enzyme, which is the primary enzyme involved in caffeine metabolism, and, experimentally, smoking decreases levels of plasma caffeine (8–12). Therefore, accelerated elimination of caffeine by tobacco smoking may lead to increased caffeine tolerance and coffee consumption (13). There may also be behavioural associations, as cigarettes and coffee are often consumed together. However, there are few data on whether smoking leads to increased coffee or caffeine consumption at the population level. Inferring causality from observational studies is difficult, due to the problems of confounding (by unmeasured or poorly measured factors) and reverse causality.

One potential way to minimise these problems is to use genetic variants which are associated with exposures as proxies for measured exposures in a Mendelian randomisation analysis (14, 15). A single nucleotide polymorphism, rs16969968, in the *CHRNA5* nicotinic receptor subunit gene influences smoking heaviness (or tobacco consumption) within smokers (16). Each additional copy of the minor allele of rs16969968 (or its proxy, rs1051730) is robustly associated with smoking, on average, one additional cigarette per day (17, 18). Of note, this variant was reported to be associated with coffee consumption in a coffee consumption GWAS, but at a nominal level of significance and in a sample combining both smokers and non-smokers (19). A recent Mendelian randomisation analysis of smoking and caffeine in a UK and a Dutch study (which did stratify into current, former and never smokers) did not find any evidence for a causal effect of smoking on caffeine intake (20). However, the sample sizes used were modest, so it is likely that these analyses were underpowered. We sought to extend this work by performing a Mendelian randomisation analysis in three large studies: the UK Biobank, the Norwegian HUNT study and the Copenhagen General Population Study.

## Subjects and Methods

### Study populations

We included individuals from three studies: the UK Biobank, which recruited over 500,000 men and women (aged 37 to 73 years) between 2006 and 2010 (21), the second wave of the Norwegian HUNT study (1995-97)(22), which invited all adults aged 20 years and older in the county of Nord-Trøndelag to participate, and the Copenhagen General Population Study (CPGS), a prospective cohort study with ongoing enrolment started in 2003 (23). Full details of the study populations, including participation flowcharts, are available in supplementary material (Supplementary Figures S1-S3).

### Genotyping

Information on genotyping of rs16969968 (UK Biobank) and rs1051730 (HUNT and CPGS) is provided in supplementary material.

### Smoking behaviour

In all studies, smoking status was classified as never smoker, former smoker, current smoker and ever smoker (current and former smokers combined). Current smokers were also asked how many cigarettes they smoked per day. Full details of how the smoking variables were derived in each study are available in supplementary material.

### Coffee and tea consumption

Coffee and tea consumption were self-reported as the number of cups consumed daily. Full details of how these variables were collected in the individual studies are available in supplementary material. Information on whether type of coffee consumption was most commonly caffeinated or decaffeinated was available in UK Biobank, but not in HUNT or Copenhagen. However, consumption of decaffeinated coffee in Norway and Denmark is low (24, 25). Information on whether tea consumption was caffeinated or decaffeinated was not available in any of the studies.

### Covariates

Analyses were adjusted for age, sex and educational attainment. A full description of the covariates used in each study is provided in supplementary material.

### Statistical analysis

Analyses were conducted in Stata (version 14. 1). Within each study, associations between smoking status (never, former and current) and smoking heaviness (cigarettes per day) and continuous measures of coffee and tea were investigated using linear regression. These analyses were adjusted for age, sex and educational attainment. Robust standard errors were used to account for non-normality of residuals as tea and coffee data tended to be right skewed.

We assessed the association between the genotypes (rs16969968/rs1051730, coded as 0,1,2 according to the number of smoking heaviness increasing alleles) and smoking heaviness using linear regression. In Mendelian randomisation analysis, we used linear regression to investigate associations of the smoking heaviness related variants with amount of coffee and tea consumption and logistic regression to investigate associations with any compared to no consumption of these drinks. These analyses were performed stratified by smoking status (never, former, current and ever). As the genetic variant is only associated with smoking heaviness in individuals who smoke, we should only see evidence for an association with coffee or tea in current smokers (and perhaps former smokers if effects are long lasting). An association with coffee or tea consumption in never smokers would suggest a direct (or pleiotropic) effect of the variant. All genetic analyses were adjusted for age, sex and principal genetic components (in UK Biobank only).

All analyses were performed within individual studies and combined in inverse variance weighted fixed effect meta-analyses. We assessed heterogeneity between associations in the different smoking categories using Cochran’s Q and the I-squared statistic. When there was evidence of heterogeneity (I-squared>50%), we used a random effects meta-analysis. Primary analyses were performed only in individuals who reported consuming coffee or tea but we also conducted sensitivity analyses including all individuals (consumers and non-consumers).

## Results

The analysis sample consisted of 114,029 individuals from UK Biobank, 56,664 individuals from the HUNT study and 78,650 individuals from the Copenhagen General Population Study (see Table 1). Smoking prevalence was higher in the HUNT study than in CGPS and UK Biobank. Coffee consumption was more common in HUNT and CGPS than in UK Biobank, but tea consumption was more common in UK Biobank than in HUNT and CGPS. Amongst consumers, individuals from UK Biobank tended to drink more cups of tea per day, but less cups of coffee per day than in HUNT and CGPS. In all studies, there was a negative correlation between coffee and tea consumption when considering individuals who consumed at least some tea or coffee (Supplementary Table S1). However, correlations were weaker in individuals reporting both tea and coffee consumption.

**Table 1.** **Characteristics of the study populations**

|  | UK Biobank (N=114,029)<sup>2</sup> |  |  | HUNT (56,664)<sup>2</sup> |  |  | CGPS (N=78,650)<sup>2</sup> |  |  |
| --- | --- | --- | --- | --- | --- | --- | --- | --- | --- |
|  | Mean/N | SD/IQR/% | Range | Mean/N | SD/IQR/% | Range | Mean/N | SD/IQR/% | Range |
| **Age: mean (SD)** | 56.9 | 7.9 | 40-73 | 49.9 | 17.1 | 19-101 | 57.6 | 13.1 | 20-99 |
| **Sex: N (%)** |  |  |  |  |  |  |  |  |  |
| Male | 53,893 | 47.3 |  | 27,021 | 47.7 |  | 35,288 | 55.1 |  |
| Female | 60,136 | 52.7 |  | 29,643 | 52.3 |  | 43,362 | 44.9 |  |
| **Educational attainment<sup>1</sup>** |  |  |  |  |  |  |  |  |  |
| Low | 20,501 | 18.1 |  | 19,759 | 36.6 |  | 17,560 | 22.4 |  |
| Middle | 40,829 | 36.1 |  | 23,514 | 43.6 |  | 27,456 | 35.0 |  |
| High | 51,687 | 45.7 |  | 10,653 | 19.8 |  | 33,376 | 42.6 |  |
| **Smoking status** |  |  |  |  |  |  |  |  |  |
| Never | 61,179 | 53.7 |  | 24,747 | 43.7 |  | 32,168 | 40.9 |  |
| Former | 39,006 | 34.2 |  | 14,350 | 25.3 |  | 31,553 | 40.1 |  |
| Current | 13,844 | 12.1 |  | 17,528 | 31.0 |  | 14,929 | 19.0 |  |
| **Number of cigarettes per day in current** | 17.0 | 8.2 |  | 11.2 | 5.7 |  | 15.4 | 9.1 |  |
| **Tea/coffee consumption** |  |  |  |  |  |  |  |  |  |
| Any tea consumption | 96,522 | 84.7 |  | 18,130 | 55.7 |  | 45,789 | 58.2 |  |
| Any coffee consumption | 89,789 | 78.7 |  | 49,991 | 91.0 |  | 70,228 | 89.3 |  |
| Any tea or coffee consumption | 111,582 | 97.9 |  | 52,512 | 95.0 |  | 77,023 | 97.9 |  |
| Tea consumption (cups per day) in tea drinkers: median (IQR) | 4 | 2,5 | 0.5-25 | 2 | 1,2 | 1-21 | 1.1 | 0.6,2.3 | 0.1-25 |
| Coffee consumption (cups per day) in coffee drinkers: median (IQR) | 2 | 1,4 | 0.5-25 | 5 | 2,5 | 1-50 | 2.9 | 1.4,4 | 0.1-25 |

1. Education defined as follows: UK Biobank (High: Degree/Professional qualifications, Middle: School/vocational qualifications, Low: No qualifications) HUNT: High: college/university education (>12 years), middle: secondary education (11-12 years), or low: primary education (<10 years). CGPS: High: University degree, Middle: Higher education (≥3 years), Low: Vocational qualifications/short education (≤3 years)/Under education/No education (School only)
<sup>2</sup>Number of participants with genotype information. N varies according to missing values on different variables.

### Associations of tea and coffee consumption and smoking with demographic factors

Similar patterns of smoking with age, sex and education were observed across the studies (see Supplementary Tables S2-S4). However, there were some differences in patterns of coffee and tea consumption by demographic factors between the studies (see Supplementary Tables S5-S7).

### Observational associations between smoking and coffee and tea consumption

In observational analyses of smoking status and smoking heaviness (cigarettes per day) with coffee and tea consumption, there was a high degree of heterogeneity between studies (I^2^ >95%). Former and current smoking were associated with higher coffee consumption (Figure 1), but there was no overall evidence from the meta-analysis that smoking status was associated with tea consumption. Amongst smokers, each additional cigarette smoked per day was positively associated with both coffee and tea consumption, although the association was larger for coffee (0.10 cups per day, 95% CI: 0.03, 0.17) than for tea (0.04 cups per day, 95% CI: -0.002, 0.07) (Figure 2). Mutual adjustment of associations for tea and coffee consumption did not make any material difference to results (see Supplementary Tables S8 and S9).

**Figure 1.**
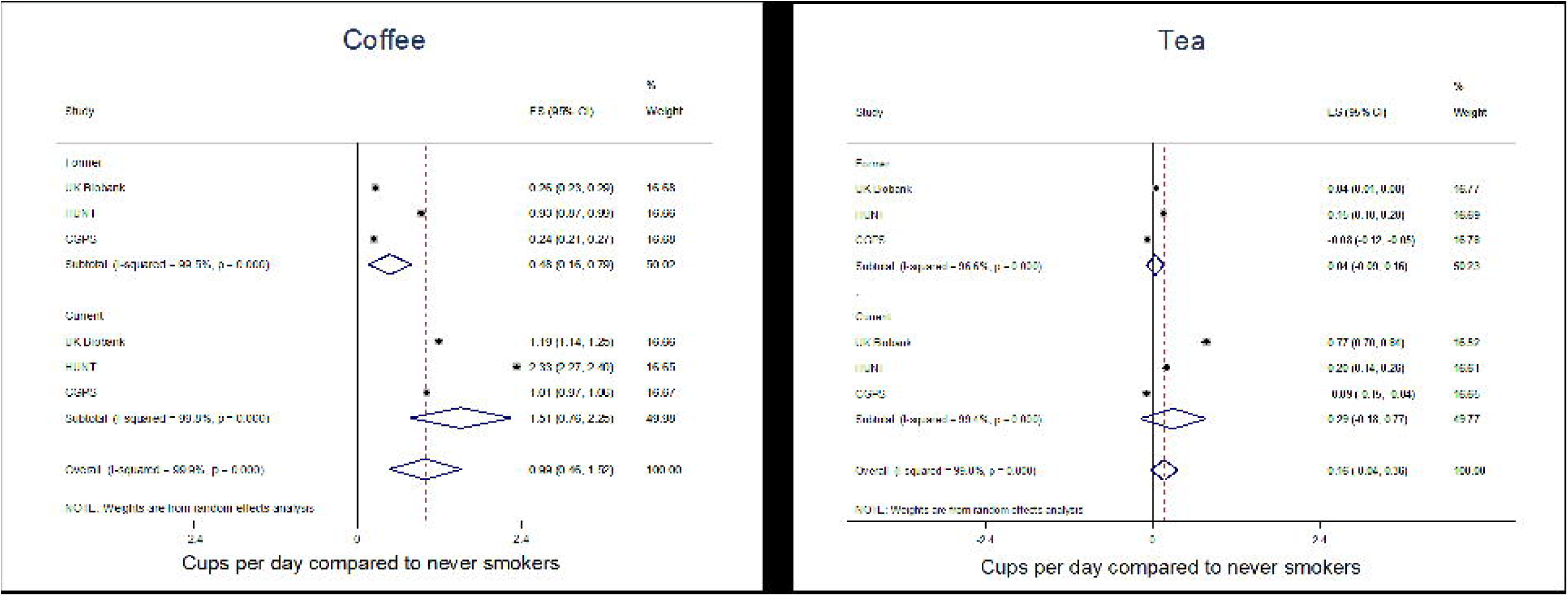
Associations between smoking status and tea and coffee consumption. Beta coefficients represent difference in coffee/tea consumption in former and current compared to never smokers. Analyses adjusted for age, sex and educational attainment. From linear regression using robust standard errors to account for non-normality of residuals. Estimates combined in a random effects meta-analysis.

**Figure 2.**
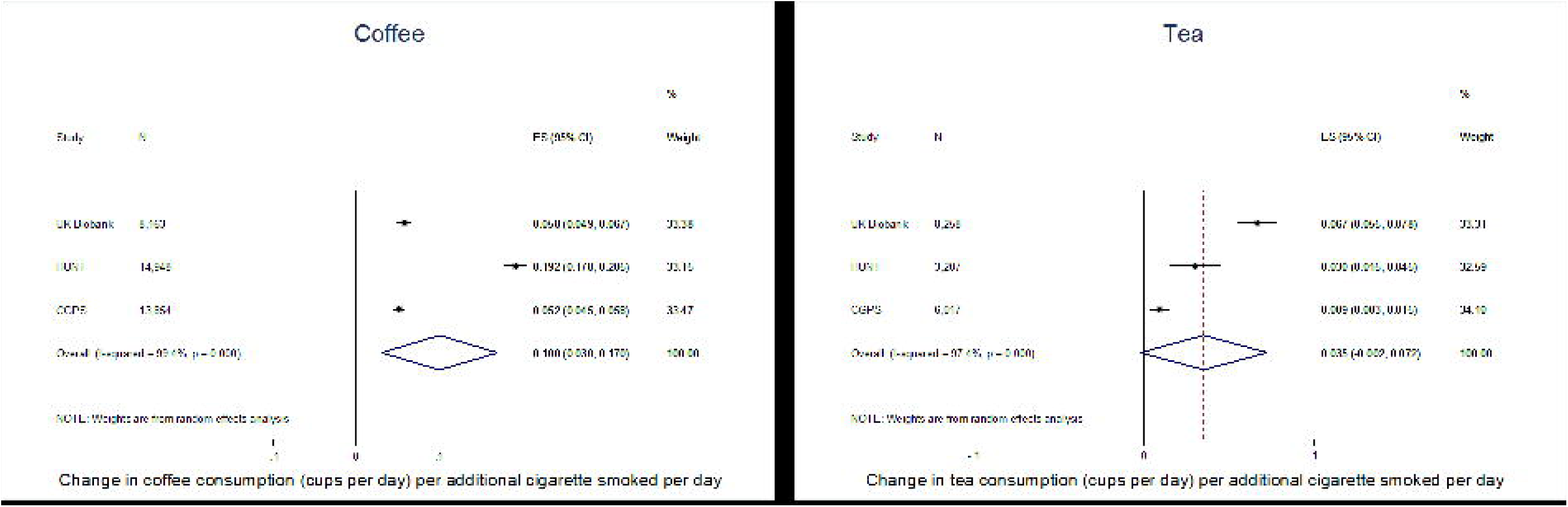
Associations between number of cigarettes per day among current smokers and coffee and tea consumption. Beta coefficients represent difference in coffee/tea consumption in current smokers per additional cigarette consumed per day. Analyses adjusted for age, sex and educational attainment. From linear regression using robust standard errors to account for non-normality of residuals. In UK Biobank, N represents number of current daily smokers only. Estimates combined in a random effects meta-analysis.

### Mendelian randomisation

In the combined estimate, each additional copy of the minor allele of rs16969968/rs1051730 was associated with smoking 0.83 additional cigarettes per day (95% CI: 0.63, 1.03, I^2^ 71%, combined using random effects meta-analysis, Supplementary Table S10). There was no clear consistent evidence that rs16969968/rs1051730 was associated with sex or education in any of the studies. When stratified by smoking status, there was some evidence in each of the studies that rs16969968/rs1051730 genotype was associated with age (age decreased per copy of the minor allele) in former and current smokers but not never smokers (see Supplementary Tables S11-S22). This is consistent with the known effects of this variant on mortality (26).

In the combined estimate from the meta-analysis, rs16969968/rs1051730 genotype was not associated with consuming any coffee compared to no coffee, or consuming any coffee or tea compared to no coffee or tea (see Supplementary Figures S4-S5). However, the minor allele was associated with reduced odds of consuming any tea compared to no tea consumption (OR: 0.94, 95% CI: 0.91, 0.97) (see Supplementary Figure S6).

There was strong evidence that the minor allele of rs16969968/rs1051730 was associated with increased coffee consumption in current smokers (Figure 3). As there was evidence for heterogeneity between studies in some smoking groups (former and ever smokers), results from random effects meta-analyses are presented. Each additional copy of the minor allele was associated with a 0.15 cup of coffee increase in consumption (95% CI: 0.11, 0.20). There was no clear evidence that rs16969968/rs1051730 was associated with coffee consumption in never smokers (beta 0.02, 95%CI: -0.003, 0.03) or in former smokers (beta 0.04, 95% CI: - 0.03, 0.10). There was evidence for heterogeneity between studies in the estimate in former smokers (I^2^ 85%). Including non-consumers of coffee in this analysis did not materially change results (Supplementary Figure S7).

**Figure 3.**
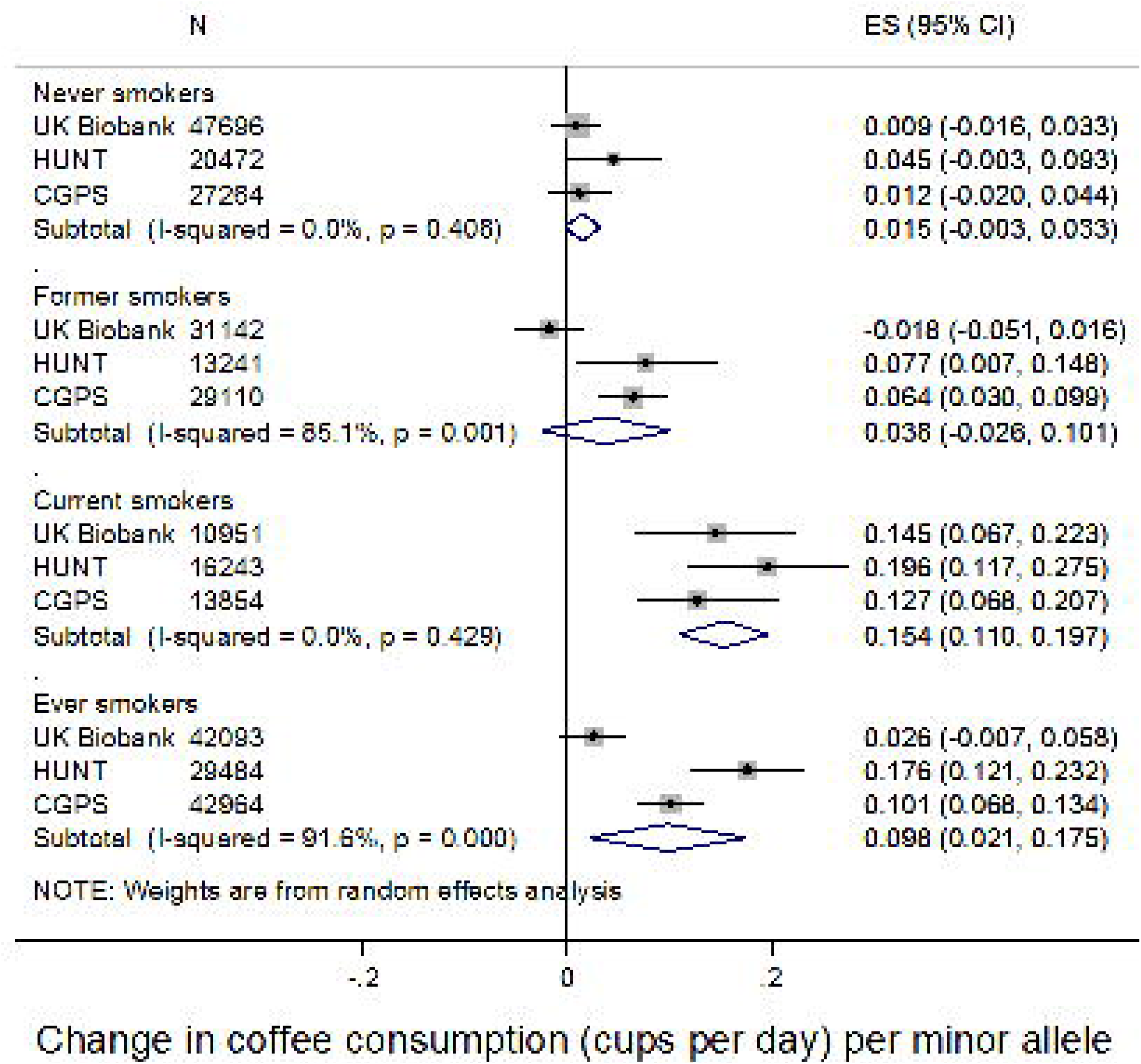
Associations between rs16969968/rs1051730 and coffee consumption. Adjusted for age, sex (in all studies) and principal components (in UK Biobank). Analyses restricted to individuals reporting at least some coffee consumption. In UK Biobank, current smokers includes daily and occasional current smokers. Estimates combined in a random effects meta-analysis.

There was no strong evidence for an association between rs16969968 and tea consumption in any of the smoking categories (Figure 4). The association was strongest in current smokers (0.04 cups per day per additional minor allele (95%CI:-0.01, 0.09)). However, there was no clear statistical evidence for differences in the association between smoking categories (P_heterogeneity_=0.22). Given the negative correlation between coffee and tea consumption, we also performed analyses combining coffee and tea consumption (by adding together cups of tea and coffee consumed). These results were similar to those observed for coffee consumption alone (see Supplementary Figure S8). Including non-consumers of tea in this analysis weakened the magnitude of association seen in both current and former smokers (Supplementary Figure S9).

**Figure 4.**
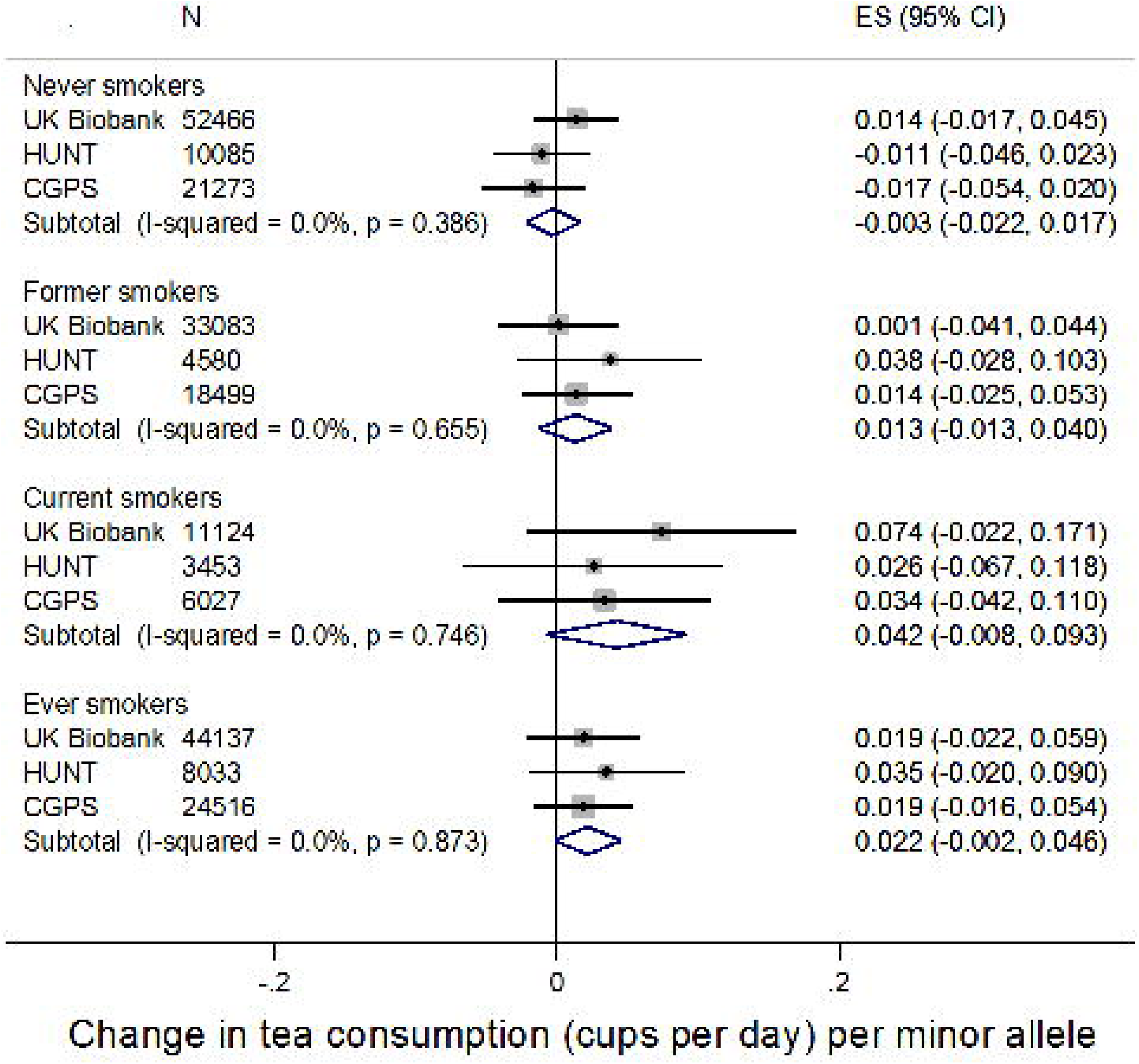
Associations between rs16969968/rs1051730 and tea consumption. Adjusted for age, sex (in all studies) and principal components (in UK Biobank). Analyses restricted to individuals reporting at least some tea consumption. In UK Biobank, current smokers includes daily and occasional current smokers. Estimates combined in a fixed effects meta-analysis.

Given the correlation between tea and coffee consumption, we performed additional analyses to investigate whether smoking might affect preference for tea or coffee. Amongst individuals consuming some tea or coffee, there was evidence that the minor allele was associated with an increase in the ratio of coffee to tea consumed amongst ever smokers (Beta 0.04, 95% CI: 0.01, 0.07) but not amongst never smokers (-0.002, 95% CI: -0.02, 0.02) (Supplementary Figure S10). Similarly, amongst individuals consuming different amounts of coffee and tea, the minor allele was associated with increased odds of drinking more coffee than tea in current (OR per minor allele: 1.06, 95% CI: 1.01, 1.10) and ever smokers (OR per minor allele: 1.02, 95% CI: 1.00, 1.05), but not never smokers (OR per minor allele: 0.99, 95% CI: 0.97, 1.01) (Supplementary Figure S11).

Within UK Biobank, data were available on type of coffee most commonly consumed. A similar pattern of results was obtained for both caffeinated and decaffeinated consumption; the minor allele was associated with higher consumption in current but not in never or former smokers (see Supplementary Figure S12). To test whether associations were limited to caffeinated beverages or reflected a general effect of smoking heaviness on thirst, we also performed a Mendelian randomisation analysis in UK Biobank with water consumption as the outcome (Supplementary Figure S13). There was no clear evidence that rs16969968/rs1051730 was associated with water consumption in any of the smoking categories, although the minor allele was associated with lower water consumption amongst current smokers, but the precision of this estimate was low.

## Discussion

In three large studies from the UK, Norway and Denmark, which have different patterns of coffee and tea consumption, we have demonstrated that smoking is positively associated with coffee intake. The results from our Mendelian randomisation analysis of coffee consumption were largely consistent across studies and provide evidence that heavier smoking causally increases coffee intake. Although the pattern of results in the Mendelian randomisation analysis was similar for tea consumption, we did not find clear evidence for a causal effect of smoking heaviness on tea consumption.

Our finding from the Mendelian randomisation analysis that the smoking increasing allele of rs16969968/rs1051730 is positively associated with coffee intake in current smokers, suggest that this association is, at least in part, due to smoking increasing coffee consumption. Lack of association in never smokers provides us with some evidence that this association is not due to pleiotropic effects of the variant (i.e., direct associations with the outcome not via the exposure of interest). Furthermore, this finding is strengthened by its consistency across studies from countries with different patterns of coffee consumption. The small positive association observed in former smokers in HUNT and CGPS suggests that there may be some residual effect that persists following smoking cessation. The reason for a lack of association in former smokers in UK Biobank is unclear; it is possible that time since quitting in UK Biobank is longer than in the other studies as the sample is restricted to older individuals. We also repeated analyses in the UK Biobank excluding occasional past smokers (as UK Biobank has a high proportion of occasional smokers) but this did not substantially alter the association.

There are two key mechanisms through which smoking may increase coffee consumption. Firstly, it is well known that cigarette smoke increases activity of the caffeine metabolising enzyme CYP1A2. Polycyclic aromatic hydrocarbons in tobacco smoke bind to the aryl hydrocarbon receptor, which activates the *CYP1A2* gene (8). Faster metabolism of caffeine in smokers (27, 28) would require them to consume more caffeine to experience its stimulating effects and avoid withdrawal, but would also allow them to consume higher levels before experiencing symptoms of caffeine toxicity (29). Secondly, the association may be due to behavioural links between smoking and coffee consumption, with smoking acting as a cue for or providing an opportunity (i.e., a cigarette break) to consume coffee (1). The association of rs16969968 with consumption of decaffeinated coffee in current smokers in UK Biobank offers some support for this behavioural explanation. It is likely that decaffeinated consumers were once caffeinated coffee consumers, so this could just reflect continuation of an established behaviour. However, as UK Biobank participants were only able to indicate their main type of coffee consumption in the questionnaire, it is likely that at least some of the coffee consumption in this “decaffeinated” analysis is caffeinated. Furthermore, the lack of clear evidence for an association with tea suggests that the coffee-smoking association is unlikely to be entirely behavioural.

Given the effect of tobacco smoke on caffeine metabolism, it is perhaps surprising that we did not find clear evidence for an association with tea consumption. It is possible that the effect of smoking is limited to coffee; the literature on the association between smoking and tea drinking is less consistent than with coffee drinking, with some studies finding positive (2) and others null or negative associations (1, 2, 30). However, this could simply reflect cultural differences in tea consumption between the countries in this analysis; in UK Biobank, which has the highest tea consumption of all the studies, there was suggestive evidence for a causal effect of smoking on tea consumption. Interestingly, the effect size in UK Biobank was around half of that observed for coffee, which might be expected given the lower caffeine content of tea (typically around half that of coffee) (2). We were also unable to distinguish between caffeinated and decaffeinated tea consumption in any of the studies; if the association is due to effects of smoking on caffeine metabolism, including decaffeinated tea would have weakened associations. Heavier smoking may cause individuals to preferentially drink coffee rather than tea due to its higher caffeine content (31). Analyses of coffee and tea preference (Supplementary Figures 8 and 9) and the negative association between the minor allele of rs16969968/rs1051730 and consumption of any tea suggest that this might be the case. Therefore, lack of association with tea consumption in this analysis may reflect lower statistical power to detect smaller associations with tea, cultural differences in tea consumption or masking of associations by preference for coffee.

There are several limitations to this analysis. Firstly the UK Biobank, had a response rate of around 5% and is not very representative of individuals of this age group in the UK. Our results could be affected by collider bias if selection into the sample is related to both coffee consumption and smoking (32). However, the consistency of the result for coffee consumption with that seen in HUNT and CGPS, which have higher response rates and are more likely to be representative of the general population, suggest that this is unlikely to be a major source of bias. Additionally the UK Biobank initial GWAS release contains about 50,000 individuals selected on the basis of smoking status and heaviness (33). However, exclusion of these individuals from the UK Biobank analysis did not materially change results (see Supplementary Figure S14). We also cannot rule out the possibility that stratification by smoking status might induce collider bias if the genetic variant itself determines smoking status (34). Whilst rs16969968/rs1051730 is largely a determinant of smoking heaviness within smokers (17) and does not appear to influence smoking initiation, there is evidence that the minor allele is associated with being a current compared to a former smoker (35). Secondly, tea and coffee consumption were self-reported and it is likely that cup size and strength of coffee differed between studies. Measurement error in our outcome would serve to decrease the precision of our findings. Finally, we have only examined one causal direction (from smoking to coffee and tea) in this analysis. Whilst we can attribute some of the association between smoking and coffee consumption to a causal effect of smoking, we cannot rule out a causal effect of caffeine or coffee on smoking behaviour or underlying genetic or environmental causes of both.

In conclusion, we provide evidence of a causal effect of smoking on coffee consumption. These findings confirm the need to consider both coffee and smoking together when considering health implications of coffee or caffeine in population studies. In addition, this may have implications for smoking cessation if individuals trying to quit smoking experience caffeine toxicity due to reduced caffeine metabolism. It has been hypothesised that overlap in symptoms between caffeine toxicity and nicotine withdrawal could increase risk of relapse to smoking (36).

